# Mitochondrial respiration contributes to the interferon gamma response in antigen presenting cells

**DOI:** 10.1101/2020.11.22.393538

**Authors:** Michael C. Kiritsy, Daniel Mott, Samuel M. Behar, Christopher M. Sassetti, Andrew J. Olive

## Abstract

The immunological synapse allows antigen presenting cells (APC) to convey a wide array of functionally distinct signals to T cells, which ultimately shape the immune response. The relative effect of stimulatory and inhibitory signals is influenced by the activation state of the APC, which is determined by an interplay between signal transduction and metabolic pathways. While toll-like receptor ligation relies on glycolytic metabolism for the proper expression of inflammatory mediators, little is known about the metabolic dependencies of other critical signals such as interferon gamma (IFNγ). Using CRISPR-Cas9, we performed a series of genome-wide knockout screens in macrophages to identify the regulators of IFNγ-inducible T cell stimulatory or inhibitory proteins MHCII, CD40, and PD-L1. Our multi-screen approach enabled us to identify novel pathways that control these functionally distinct markers. Further integration of these screening data implicated complex I of the mitochondrial respiratory chain in the expression of all three markers, and by extension the IFNγ signaling pathway. We report that the IFNγ response requires mitochondrial respiration, and APCs are unable to activate T cells upon genetic or chemical inhibition of complex I. These findings suggest a dichotomous metabolic dependency between IFNγ and toll-like receptor signaling, implicating mitochondrial function as a fulcrum of innate immunity.

## Introduction

During the initiation of an adaptive immune response, the antigen presenting cell (APC) serves as an integration point where tissue-derived signals are conveyed to T cells. Myeloid APCs, such as macrophages and dendritic cells (DCs), are responsible for the display of specific peptides in complex with MHC molecules, and for the expression of co-signaling factors that tune the T cell response (1). The expression of stimulatory or inhibitory co-signaling molecules depends on the local immune environment and activation state of the APC (2). In particular, interferon gamma (IFNγ) stimulates the surface expression of MHC proteins (3–9), co-stimulatory proteins such as CD40, and the secretion of cytokines like IL-12 and IL-18 (10), to promote T cell activation and the production of IFNγ-producing T-helper type 1 (Th1) effector cells (11–15). In the context of local inflammation, pattern recognition receptor (PRR) ligands and endogenous immune activators can collaborate with IFNγ to induce the expression of co-inhibitory molecules, like programmed death-ligand 1 (PD-L1) (16–22), which ligates T cell programmed death receptor 1 (PD1) to limit immune activation and mitigate T cell-mediated tissue damage (23–26).

IFNγ mediates these complex effects via binding to a heterodimeric surface receptor (27–30). The subunits of the complex, IFNGR1 and IFNGR2, assemble once IFNGR1 is bound by its ligand (31, 32). Complex assembly promotes the phosphorylation of janus kinases 1 and 2 (JAK1 and JAK2) followed by activation of the signal transducer and activation of transcription 1 (STAT1) (33). Phosphorylated STAT1 then dimerizes and translocates to the nucleus to activate the transcription of genes containing promoters with IFNγ-activated sequences (GAS), which includes other transcription factors such as interferon regulatory factor 1 (*Irf1*) that amplify the expression of a large regulon that includes T cell co-signaling molecules (34, 35). The importance of this signaling pathway is evident in a variety of diseases including cancer (36–40), autoimmunity (41, 42), and infection (43). Individuals with inborn deficiencies in IFNγ signaling, including mutations to the receptor (44, 45), suffer from a defect in Th1 immunity that results in an immunodeficiency termed Mendelian susceptibility to mycobacterial disease (MSMD) (46–49). Conversely, antagonists of IFNγ-inducible inhibitory molecules, such as PD-L1, are the basis for checkpoint inhibitor therapies that effectively promote T cell-mediated tumor destruction (26, 28, 50–55). While the obligate components of the IFNγ signaling pathway are well known, characterization of additional regulators of this response promises to identify both additional causes of immune dysfunction and new therapeutic targets.

Recent data suggests that cellular metabolism is an important modulator of the APC-T cell interaction. In particular, microbial stimulation of PRR receptors on the APC induces glycolytic metabolism and this shift in catabolic activity is essential for cellular activation, migration, and CD4+ and CD8+ T cell activation (18, 56–70). The metabolic state of the T cell is also influenced by the local environment and determines both effector function and long-term differentiation into memory cells (71, 72). Like PRR signaling, IFNγ stimulation has been reported to stimulate glycolysis and modulate cellular metabolism in macrophages (66, 73). However, the effects of different metabolic states on IFNγ-stimulated APC function remains unclear.

To globally understand the cellular pathways that influence IFNγ-dependent APC function, we used a CRISPR-Cas9 knockout library (74) in macrophages to perform a series of parallel forward-genetic screens for regulators of three IFNγ-inducible co-signaling molecules: MHCII, CD40, and PD-L1. We identified positive and negative regulators that controlled each marker, underscoring the complex regulatory networks that influence the interactions between APCs and T cells. Pooled analysis of the screens uncovered shared regulators that contribute to the global IFNγ response. Prominent among these general regulators was complex I of the respiratory chain. We report that the activity of the IFNγ receptor complex and subsequent transcriptional activation depends on mitochondrial function in both mouse and human myeloid cells. Experimental perturbation of respiration inhibits the capacity of both macrophages and dendritic cells to stimulate T cells, identifying mitochondrial function as a central point where local signals are integrated to determine APC function.

## Results

### Forward genetic screen identifies regulators of IFNγ-inducible MHCII, CD40 and PD-L1 cell surface expression

To investigate the diverse regulatory pathways underlying the IFNγ response, we examined the expression of three functionally distinct cell surface markers that are induced by IFNγ. Stimulation of Cas9-expressing immortalized bone marrow-derived macrophages with IFNγ for 24 hours resulted in the upregulation of T cell stimulatory molecules, major histocompatibility complex class II (MHCII) and CD40, and the inhibitory ligand PD-L1 (*Cd274*), on the cell surface (Figure 1A). To identify genes that regulate the expression of these markers, Cas9-expressing macrophages were transduced with a lentiviral genome-wide knockout (KO) library containing four single guide RNAs (sgRNAs) per protein-coding gene and 1000 non-targeting control (NTC) sgRNAs (74). The knockout library was then stimulated with IFNγ, and fluorescently activated cell sorting (FACS) was used to select for mutants with high or low cell surface expression of each individual marker (Figure 1B). For each of the three surface markers, positive and negative selections were performed in duplicate. The sgRNAs contained in the input library and each sorted population were amplified and sequenced (Figure 1A,B).

**Figure 1.**
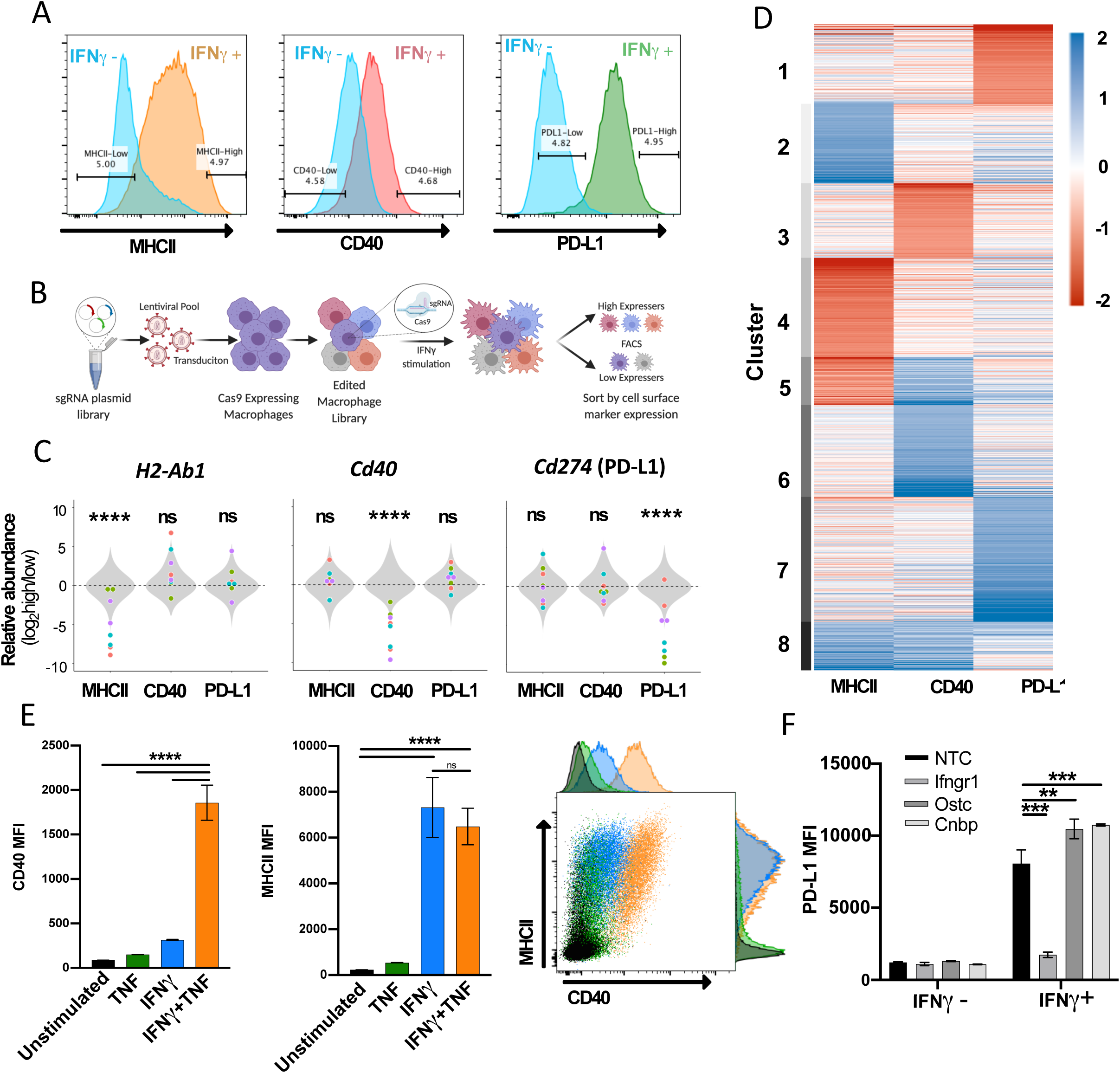
Forward genetic screen to identify regulators of the IFNγ response. A) Representative histograms of the three selected cell surface markers targeted in macrophage CRISPR screens: MHCII, CD40, and PD-L1. Blue histograms indicate expression of each marker in unstimulated macrophages and alternatively colored histograms show expression following 24 hour stimulation with recombinant murine IFNγ (10ng/mL). Gates used for sorting “high” and “low” populations are shown. B) Schematic of CRISPR screens. C) Relative enrichment of select positive control (points) and all 1000 non-targeting control sgRNAs (gray distribution) are plotted as a function of their log2 fold enrichment (“high” vs “low” bins). Data are from both replicate selections for each sgRNA (sgRNA denoted by shape). D) Heatmap of β scores from CRISPR analysis, ordered according to k-means clustering (k=8) of the 5% most enriched or depleted genes in each screen. E) Macrophages were stimulated for 24 hours with TNF (25ng/mL), IFNγ (10ng/mL) or both TNF and IFNγ. Mean fluorescence intensity (MFI) of CD40 and MHCII were quantified by flow cytometry. Data are mean ± the standard deviation for 3 biological replicates. Representative scatter plot from two independent experiments is provided. F) Macrophages transduced with sgRNA targeting *Stat1, Ostc, Cnbp*, or a NTC control were cultured with or without IFNγ for 24 hours and cell surface expression of PD-L1 (MFI) was quantified by flow cytometry. For each genotype, data are the mean of cell lines with two independent sgRNAs ± the standard deviation. Data are representative of three independent experiments. Statistical testing in panel C was performed with Tukey’s multiple comparisons test. Within each screen, the sgRNA effects for each gene were compared to the distribution non-targeting control sgRNAs. Statistical testing in panels E and F was performed by one-way ANOVA with Holm-Sidak multiple comparisons correction. p values of 0.05, 0.01, 0.001, and 0.001are indicated by *, **, ***, and ****

To estimate the strength of selection on individual mutant cells, we specifically assessed the relative abundance of cells harboring sgRNAs that target each of the surface markers that were the basis for cell sorting. When the abundances of sgRNAs specific for *H2-Ab1* (encoding the MHCII, H2-I-A beta chain), *Cd40*, or *Cd274* (PD-L1) were compared between high- and low-expressing cell populations, we found that each of these sgRNAs were significantly depleted from the cell populations expressing the targeted surface molecule, while each had no consistent effect on the expression of non-targeted genes (Figure 1C). While not all individual sgRNAs produced an identical effect, we found that targeting the genes that served as the basis of sorting altered the mean relative abundance 30-60 fold, demonstrating that all selections efficiently differentiated responsive from non-responsive cells.

We next tested for statistical enrichment of sgRNAs using MAGeCK-MLE (75), which employs a generalized linear model to identify genes, and by extension regulatory mechanisms, controlling the expression of each surface marker. This analysis correctly identified the differential representation of sgRNAs targeting genes for the respective surface marker in the sorted populations in each screen, which were found in the top 20 ranked negative selection scores (Ranks: *H2-Ab1* = 20, *Cd40* = 1, *Cd274* = 3; Table S1). Upon unsupervised clustering of β scores for the most highly enriched genes in each screen (top 5%, positive or negative) both common and pathway-specific effects were apparent (Figure 1D; Table S2). A small number of genes assigned to Cluster 1, including the IFNγ receptor components (*Ifngr1* and *Ifngr2)*, were strongly selected in the non-responsive population in all three selections. However, many mutations appeared to preferentially affect the expression of individual surface markers, including a number of known pathway-specific functions. For example, genes previously shown to specifically control MHCII transcription, such as *Ciita*, *Rfx5*, *Rfxap*, *Rfxank*, and *Creb1* (8, 76–78) were found in Cluster 4 along with several novel regulators that appear to be specifically required for this pathway. MHCII-specific factors are reported in an accompanying study (79).

Genes specifically required for CD40 expression in Cluster 3 included the heterodimeric receptor for TNF. *Tnfrsf1a* and *Tnfrsf1b* were the 6^th^ and 50^th^ lowest β scores in the CD40 screen, respectively. Previous studies suggested that TNF stimulation enhances IFNγ-mediated CD40 expression in hematopoietic progenitors (80), and we confirmed this observation in macrophages (Figure 1E). We observed a 6-fold higher induction of CD40 in macrophages stimulated with a combination of IFNγ and TNF compared to IFNγ alone. This synergy was specific to CD40 induction, as we did not observe any enhancement of IFNγ-induced MHCII expression by TNF addition.

Several recent studies identified genes that control PD-L1 expression in cancer cell lines(28, 53, 55, 81–86), and we validated the PD-L1-associated clusters using these candidates. Our analysis found the previously-described negative regulators, *Irf2* (87), *Keap1*, and *Cul3* (88–90) in the PD-L1-related Cluster 7, along with novel putative negative regulators such as the oligosaccharlytransferase complex subunit *Ostc* and the transcriptional regulator, *Cnbp*. We generated knockout macrophages for each of these novel candidates and confirmed that mutation of these genes enhances the IFNγ-dependent induction of PD-L1 surface levels (Figure 1F). Cumulatively, these data delineate the complex regulatory networks that shape the IFNγ response.

### **M**itochondrial complex I is a positive regulator of the IFNγ response

To identify global regulators of the IFNγ response, we performed a combined analysis, reasoning that treating each independent selection as a replicate measurement would increase our power to identify novel pathways. We used MAGeCK to calculate a selection coefficient (β) for each gene by maximum likelihood estimation (75). By combining the 24 available measurements for each gene (three different markers, each selection in duplicate, and four sgRNAs per gene), we found that the resulting selection coefficient reflected the global importance of a gene for the IFNγ response (Table S3). The most important positive regulators corresponded to the proximal IFNγ signaling complex (Figure 2A). Similarly, we identified known negative regulators of IFNγ signaling, including the protein inhibitor of activated Stat1 (*Pias1*) (91), protein tyrosine phosphatase non-receptor type 2 (*Ptpn2*) (84), Mitogen activate protein kinase 1 (*Mapk1)*, and suppressor of cytokine signaling 1 (*Socs1*) and 3 (*Socs3*).

**Figure 2.**
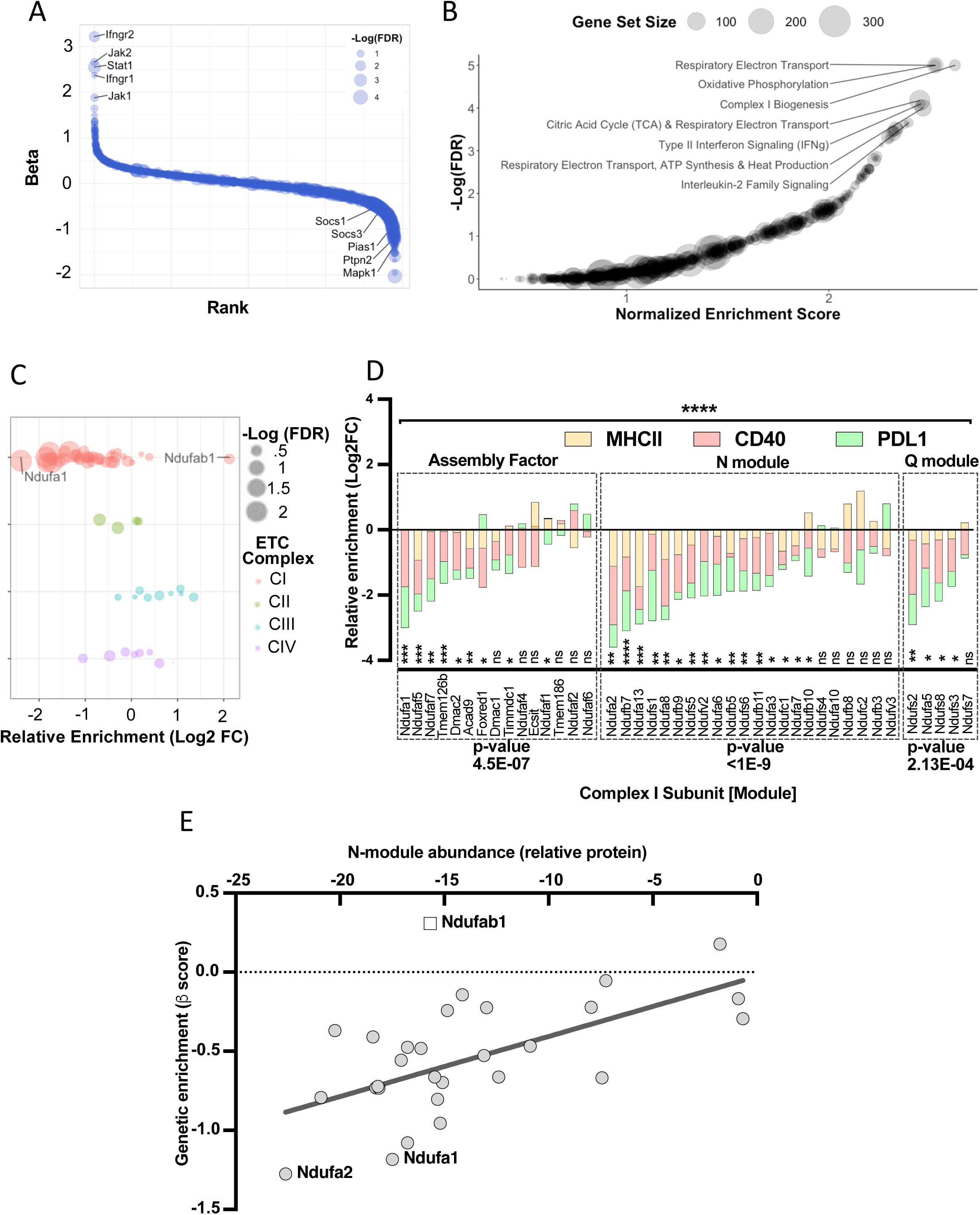
Global analysis of knockout libraries implicates mitochondrial complex I is a positive regulator of the IFNγ response. A) Rank plot of the combined analysis for all genome-wide knockout screens. Gene ranks (x-axis) were determined by maximal likelihood estimation (MLE). Known positive (left) and negative (right) regulators of IFNγ-mediated signaling are highlighted. The q-value (false discovery rate) for each gene is indicated by dot size (-Log_10_ FDR). B). Gene set enrichment analysis (GSEA) is based on the ranked list of positive regulators. Non-redundant pathways with a normalized enrichment score (NES) exceeding 2.0 and a false discovery rate (FDR) below 0.025 are labeled. C) Relative enrichment (log2 fold change between “high” and “low” bins) of genes which comprise the mitochondrial respirasome (GeneOntology 0005746) and were targeted in the CRISPR KO library. Respirasome components are grouped by ETC complex. FDR is based on MAGeCK-MLE. D) Screen-specific enrichment score is plotted for Complex I structural subunits and assembly factors. The statistical enrichment of a gene (e.g. *Ndufa1*) or module (e.g. N) was calculated using a binomial distribution function to calculate the probability that observed sgRNAs under examination would be depleted or enriched given the expected median probability. P values of 0.05, 0.01, 0.001, and 0.001are indicated by *, **, ***, and ****. E) Correlation between the relative effect of each complex I subunit on the structural integrity of the N-module (x-axis) with the relative requirement of each complex I subunit for the IFNγ-response (y-axis; β score, as in Panel D). The Pearson correlation coefficient (r) was calculated to be 0.6452 (95% confidence interval 0.3584 to 0.8207; p-value = 0.0002. As *Ndufab1* (empty square) is an essential gene, its detection in the library indicates editing did not eliminate function; therefore, it was excluded from correlation analysis.

We performed gene set enrichment analysis (GSEA) using a ranked list of positive regulators from the combined analysis (Table S4) (92). Among the top enriched pathways was a gene set associated with type II interferon (e.g., IFNγ) signaling (normalized enrichment score = 2.45, q-value = 7.98e-5), validating the approach. GSEA identified a similarly robust enrichment for gene sets related to mitochondrial respiration and oxidative phosphorylation (Figure 2B). In particular, we found a significant enrichment of gene sets dedicated to the assembly and function of the NADH:ubiquinone oxidoreductose (hereafter, “complex I”) of the mitochondrial respiratory chain. Complex I couples electron transport with NADH oxidation and is one of four protein complexes that comprise the electron transport chain (ETC) that generates the electrochemical gradient for ATP biosynthesis. To confirm the GSEA results, we examined the combined dataset for individual genes that make up each complex of the ETC (Figure 2C). This analysis demonstrated that sgRNAs targeting components of complexes II, III or IV had minimal effects on the expression of the IFNγ-inducible surface markers tested. In contrast, the disruption of almost every subunit of complex I impaired the response to IFNγ, with the notable exception of *Ndufab1*. As this gene is essential for viability (93), we assume that cells carrying *Ndufab1* sgRNAs retain functional target protein.

To investigate the contribution of specific complex I components to different IFNγ- stimulated phenotypes, we reviewed the surface marker-specific enrichment scores for genes that contribute to the complex assembly, the electron-accepting N-module, or the electron-donating Q module (93–98). Of the 48 individual assembly factors or structural subunits of complex I present in our mutant library, 29 were significantly enriched as positive regulators in the global analysis and were generally required for the induction of all IFNγ-inducible markers (Fig. 2D). The enrichment for each functional module in non-responsive cells was statistically significant. However, not all individual complex I components were equally enriched, which could reflect either differential editing efficiency or distinct impacts on function. To investigate the latter hypothesis, we compared our genetic data with a previous proteomic study that quantified the effect of individual complex I subunits on the stability of the largest subcomplex, the N-module (93). For a given subunit, we found a significant correlation between the magnitude of enrichment in our genetic screen and its effect on the structural stability of the module (Fig. 2E), specifically implicating the activity of complex I in the IFNγ response.

To directly test the predictions of the screening data, we used CRISPR to generate individual macrophage lines that were deficient for complex I subunits. We first validated the expected metabolic effects of complex I disruption by comparing the intracellular ATP levels in macrophages carrying non-targeting control sgRNA (sgNTC) with sg*Ndufa1* and sg*Ndufa2* lines. When cultured in media containing the glycolytic substrate, glucose, all cell lines produced equivalent amounts of ATP (Figure 3A). However, when pyruvate was provided as the sole carbon source, and ATP generation depends entirely upon flux through ETC and oxidative phosphorylation (OXPHOS), both sg*Ndufa1* and sg*Ndufa2* macrophages contained decreased ATP levels compared to sgNTC cells (Figure 3B). To confirm the glycolytic dependency of complex I mutant macrophages, we grew cells in complete media with glucose and treated with the ATP synthase (complex V) inhibitor, oligomycin, which blocks ATP generation by OXPHOS. While oligomycin reduced ATP levels in sgNTC macrophages, this treatment had no effect in sg*Ndufa1* and sg*Ndufa2* cells (Supplementary Figure 1A), confirming that these complex I-deficient cells rely on glycolysis for energy generation. IFNγ treatment slightly reduced ATP levels in glucose containing media but did not differentially affect cell lines (Figure 3A). Throughout these experiments we found that the sg*Ndufa1* mutant showed a greater OXPHOS deficiency than the sg*Ndufa2* line.

**Figure 3.**
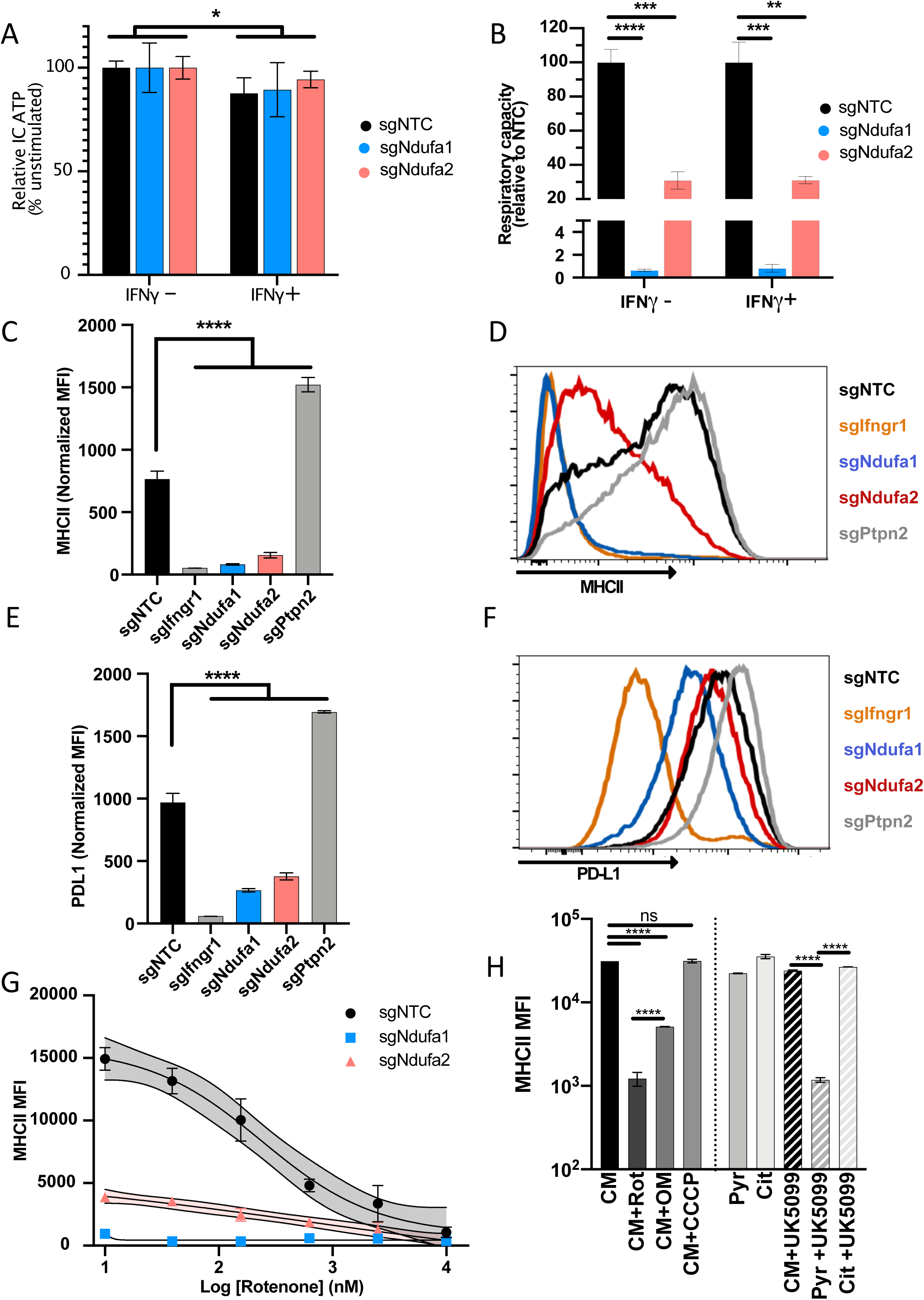
Complex I is necessary for IFNγ-induced MHCII and PD-L1 expression. Metabolic phenotypes in macrophage mutants were confirmed using ATP abundance following culture in media containing only (A) glucose or (B) pyruvate. Values are normalized to the average respiratory capacity of non-targeting control macrophages (NTC) and are the mean ± the standard deviation for 4 biological replicates. Statistical testing within each condition (with or without IFNγ for 24h) was performed by one-way ANOVA with Dunnett’s multiple comparisons correction. (C-F) Non-targeting control (NTC), positive control (sg*Ifngr1* and sg*Ptpn2*) and complex I mutant (sg*Ndufa1* and sg*Ndufa2*) macrophages were stimulated for 24 hours with recombinant murine IFNγ. Plotted values in C and E are the geometric mean fluorescence intensity (MFI) for a given mutant normalized to an internal control present in each well; for each gene, the data are the mean for two independent sgRNAs ± the standard deviation. Representative histograms are provided in D and F. Data are representative of >5 independent experiments. G) MHCII MFI of macrophages stimulated with IFNγ and treated with rotenone at the indicated concentrations for 24 hours. Mean ± the standard deviation for 2 biological replicates are shown. Data are representative of four independent experiments. H) Left: MHCII MFI on macrophages cultured in complete media (CM) and stimulated with IFNγ and the indicated inhibitors for 24 hours. Right: MHCII MFI on macrophages cultured in CM or media containing only pyruvate (Pyr) or citrate (Cit) with or without UK5099 and stimulated with IFNγ for 24 hours. Mean ± standard deviation for 2 or 3 biological replicates is indicated. Data are representative of four independent experiments. Statistical testing was performed by one-way ANOVA with Tukey correction for multiple hypothesis testing. p values of 0.05, 0.01, 0.001, and 0.001are indicated by *, **, ***, and ****.

We next compared the response to IFNγ in macrophages lacking *Ndufa1* and *Ndufa2* with those carrying CRISPR-edited alleles of *Ifngr1* or the negative regulator of signaling, *Ptpn2*. As CD40 was found to rely on more complex inputs for expression, which include TNF (Figure 1E), we relied on MHCII and PD-L1 as markers of the IFNγ response for subsequent studies. As expected, and consistent with the genetic screen, we found that the loss of *Ifngr1* or *Ptpn2* either abrogated or enhanced the response to IFNγ, respectively. Also consistent with predictions, mutation of complex I genes significantly reduced the IFNγ-dependent induction of MHCII and PD-L1 compared to sgNTC (Figure 3C-F). The *Ndufa1* mutation that abrogates OXPHOS, reduced MHCII induction to the same level as *Ifngr1*-deficient cells. To confirm these results using an orthologous method we treated cells with the complex I inhibitor, rotenone (99). This treatment caused a dose-dependent inhibition of the IFNγ-induced MHCII expression in sgNTC macrophages (Figure 3G) and had a similar inhibitory effect on the residual IFNγ response in *Ndufa2*-deficient cells. Together these results confirm that complex I is required for the induction of immunomodulatory surface molecules in response to IFNγ.

To investigate what aspect of mitochondrial respiration contributes to the IFNγ response, we inhibited different components of the ETC. All inhibitors were used at a concentration that abrogated OXPHOS-dependent ATP generation (Supplementary Figure 1B). The complex V inhibitor, oligomycin, inhibited the IFNγ-induced MHCII expression, albeit to a lesser extent than direct complex I inhibition with rotenone (Figure 3H). This partial effect could reflect an inability to dissipate the proton motive force (PMF), which inhibits electron flux throughout the ETC, including through complex I (100). Carbonyl cyanide m-chlorophenyl hydrazone (CCCP) disrupts mitochondrial membrane potential and OXPHOS while preserving electron flux. CCCP had no effect on the IFNγ response, indicating that ATP generation is dispensable for IFNγ responsiveness and highlighting a specific role for complex I activity.

We then altered the media composition to test the sufficiency of mitochondrial respiration to drive IFNγ responses independently from aerobic glycolysis. IFNγ was found to stimulate MHCII expression to a similar degree in macrophages cultured in complete media with glucose as in media containing only pyruvate or citrate, which must be catabolized via mitochondrial respiration (Figure 3H). Inhibition of mitochondrial pyruvate import with the chemical inhibitor, UK5099 (101), abrogated MHCII induction in cultures grown in pyruvate, but not in citrate, which is imported via a UK5099-independent mechanism. Taken together these results suggest that cellular respiration is both necessary and sufficient for maximal expression of the IFNγ- inducible surface markers MHCII and PD-L1.

### Mitochondrial function is specifically required IFNγ-dependent responses

The mitochondrial-dependency of the IFNγ response contrasted with the known glycolytic-dependency of Toll-like receptor (TLR) signaling, suggesting that TLR responses would remain intact when complex I was inhibited. Indeed, not only were TLR responses intact in sg*Ndufa1* and sg*Ndufa2* mutant macrophages, these cells secreted larger amounts of TNF or IL-6 than sgNTC cells in response to the TLR2 ligand, Pam3CSK4. (Figure 4A). Thus, the glycolytic dependency of these cells enhanced the TLR2 response, indicating opposing metabolic dependencies for IFNγ and TLR signaling.

**Figure 4.**
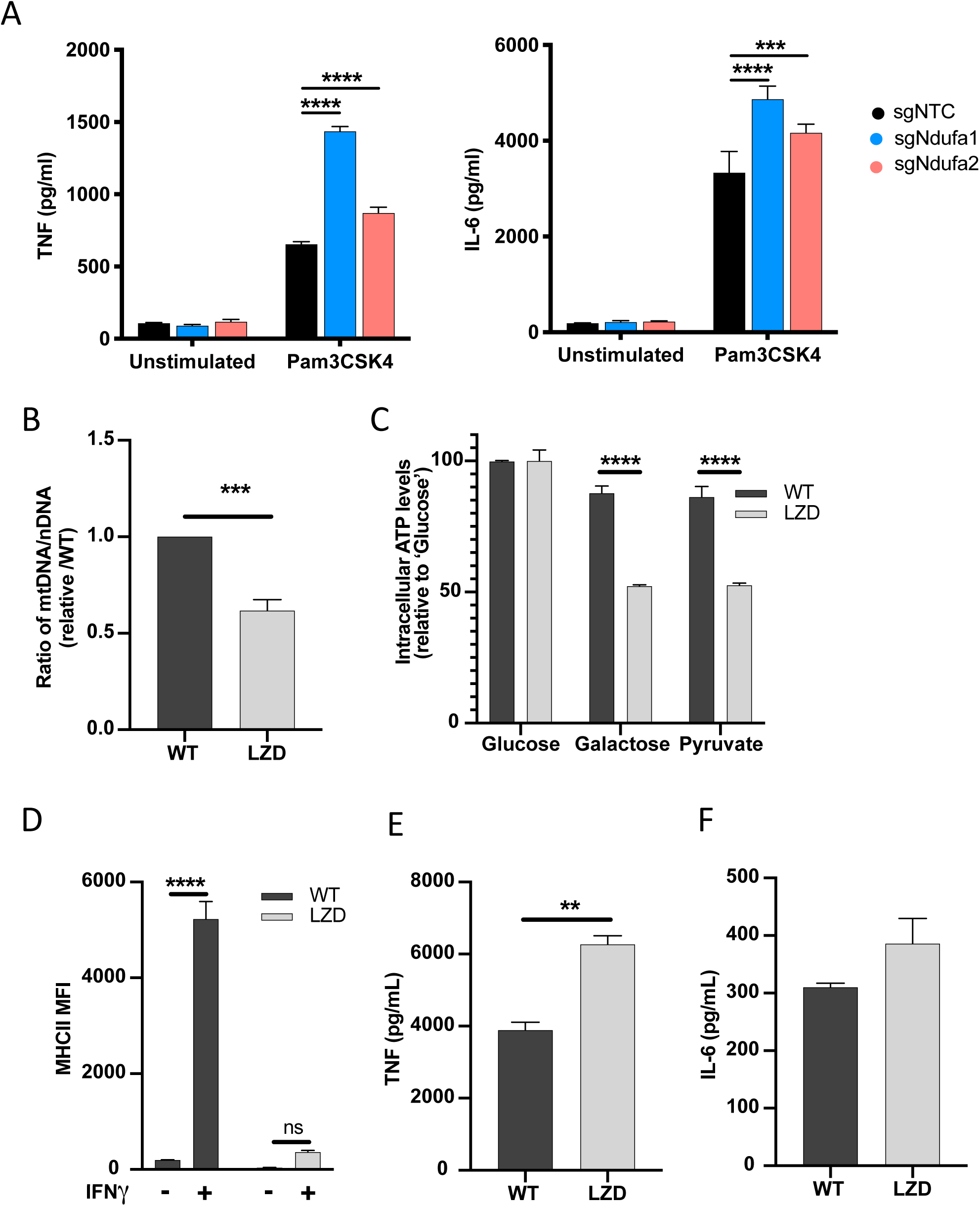
Diminished mitochondrial function specifically limits IFNγ-dependent responses. A) TNF and IL-6 production by NTC or complex I mutant macrophages stimulated with Pam3CSK4 for 24 hours was determined by ELISA. Statistical testing between mutant and NTC macrophages from triplicate samples was performed by ANOVA with Dunnett’s correction for multiple comparisons. Data are representative of two independent experiments. B) qPCR determination of relative mitochondrial genomes present per nuclear genome in macrophages cultured in vehicle (WT) or 50 ug/mL linezolid (LZD). C_t_ values were normalized to reference nuclear gene hexokinase 2 (*Hk2*) and plotted as abundance relative to WT. Data were analyzed by two-way unpaired t-test. C) ATP abundance in control or LZD-conditioned macrophages cultured in 10mM glucose, galactose or pyruvate. ATP values normalized to mean of 10mM glucose and plotted as percent. Mean ± the standard deviation for 2 biological replicates of each condition. Differences were tested by two-way ANOVA using the Sidak method to correct for multiple hypothesis testing. D) MFI of MHCII was determined by flow cytometry on control or LZD-conditioned macrophages following 24 hour stimulation with IFNγ. Mean ± the standard deviation for 2 biological replicates of each condition and representative of two independent experiments. Differences were tested by two-way ANOVA using the Tukey method to correct for multiple hypothesis testing. E and F) Secretion of TNF and IL-6 in WT and LZD-conditioned macrophages following Pam3CSK4 stimulation for 6 hours was quantified by ELISA. Mean ± the standard deviation for 3 biological replicates of each condition and two independent experiments. Data were analyzed by two-way unpaired t-test. p values of 0.05, 0.01, 0.001, and 0.001are indicated by *, **, ***, and ****.

Whether the effects of complex I on macrophage responsiveness was the result of reduced mitochondrial respiratory function or secondary to cellular stress responses, such as radical generation, remained unclear. To more directly relate mitochondrial function to these signaling pathways, we created cell lines with reduced mitochondrial mass. Macrophages were continuously cultured in linezolid (LZD), an oxazolidinone antibiotic that inhibits the mitochondrial ribosome (102–104). This treatment produced a cell line with ∼50% fewer mitochondrial genomes per nuclear genome and a corresponding decrease in OXPHOS capacity, compared to control cells grown in the absence of LZD (Figure 4B,C). Cells were cultured without LZD for 16 hours and then stimulated with either IFNγ or Pam3CSK4. Consistent with our complex I inhibition studies, we found this reduction in mitochondrial mass nearly abrogated the IFNγ-dependent induction of MHCII (Figure 4D), while the TLR2-dependent secretion of TNF and IL-6 was preserved or enhanced (Figure 4E and 4F). Thus, mitochondrial activity, itself, is necessary for a robust IFNγ response.

To further address potential secondary effects of mitochondrial inhibition on the IFNγ response, we investigated the role of known oxygen or nitrogen radical-dependent regulators (Supplementary Figure 1C-G). Inhibition of ROS generation by replacing glucose with galactose (66, 100, 105) had no effect on IFNγ-induced MHCII induction. Similarly, neutralization of cytosolic or mitochondrial radicals with N-acetylcysteine or MitoTempo, respectively, had no effect on MHCII induction either alone or in combination with ETC inhibition. The role of the cytosolic redox sensor, *HIF1α* (106, 107) was addressed by chemically stabilizing this factor with dimethyloxalylglycine (DMOG). A potential role for nitric oxide production was addressed with the specific NOS2 inhibitor 1400W (60, 66, 108). Neither of these treatments affected IFNγ-induced MHCII cell surface expression in the presence or absence of simultaneous Pam3CSK4, further supporting a direct relationship between mitochondrial respiratory capacity and the IFNγ response.

### Complex I is specifically required for IFNγ signaling in human cells

To understand the function of complex I during IFNγ-stimulation in human cells, we used monocyte-derived macrophages (MDM) from peripheral blood of healthy donors. As in our mouse studies, we assessed the response of these cells to IFNγ or Pam3CSK4 by quantifying the abundance of IFNγ-inducible surface markers or cytokines that were optimized for human cells. Since HLA-DR is not strongly induced by IFNγ, we included ICAM1 in addition to CD40 and PD-L1 as surface markers. As seen in the murine model, rotenone inhibited the IFNγ-mediated induction of all three markers (Figure 5A). TLR2 responses were assessed by the production of TNF and IL-1β. Upon Pam3CSK4 stimulation, rotenone significantly enhanced the secretion of IL-1β and TNF (Figure 5B). While simultaneous treatment with both IFNγ and Pam3CSK4 produced the previously described inhibition of IL-1β (109), rotenone still did not decrease the production of these TLR2 dependent cytokines. Thus, as we observed in mouse cells, complex I is specifically required for IFNγ signaling in human macrophages.

**Figure 5.**
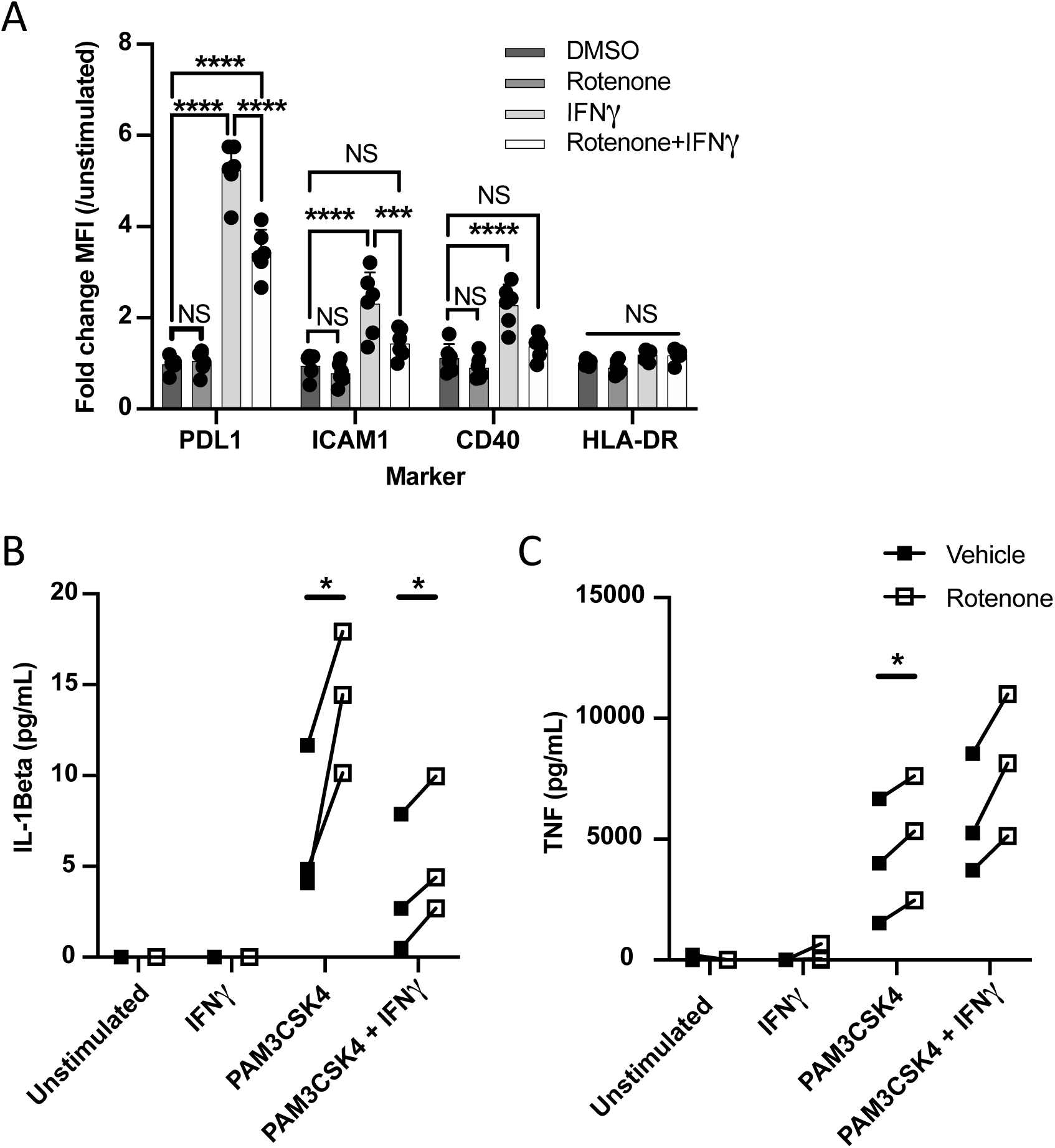
Complex I is specifically required for IFNγ signaling in human cells. A) CD14+ monocytes from healthy human donors were differentiated into macrophages. MFI of cell surface markers PD-L1, ICAM1, CD40 and HLA-DR was determined by flow cytometry following stimulation with IFNγ and/or inhibition of complex I with rotenone (10uM) for 24 hours. Data are representative of two independent experiments and values are normalized to donor-specific unstimulated/vehicle control. Mean ± the standard deviation for 6 biological replicates of each condition. Differences were tested by two-way ANOVA using the Sidak-Holm method to correct for multiple hypothesis testing. B and C) Quantification of IL-1B and TNF production from primary human macrophages, measured by ELISA from cell supernatants following stimulation. Lines connect values for individual donors treated with vehicle (DMSO, black squares) or rotenone (empty squares). Differences were tested by repeat-measure two-way ANOVA using the Sidak-Holm method to correct for multiple hypothesis testing. p values of 0.05, 0.01, 0.001, and 0.001are indicated by *, **, ***, and ****.

### Complex I inhibition reduces IFNγ receptor activity

To understand how complex I activity was shaping the IFNγ response, we first determined whether its effect was transcriptional or post-transcriptional by simultaneously monitoring mRNA and protein abundance over time. Surface expression of PD-L1 was compared with the gene’s mRNA abundance, while the surface expression of MHCII was compared with the mRNA abundance of *Ciita*, the activator of MHCII expression that is initially induced by IFNγ (Figure 6 A,B). In both cases, mRNA induction preceded surface expression of the respective protein. More importantly, both mRNA and protein expression of each marker was diminished to a similar degree in sg*Ndufa1* and sg*Ndufa2*, compared to sgNTC cells. Thus, a deficit in transcriptional induction could account for the subsequent decrease in surface expression observed in complex I deficient cells.

**Figure 6.**
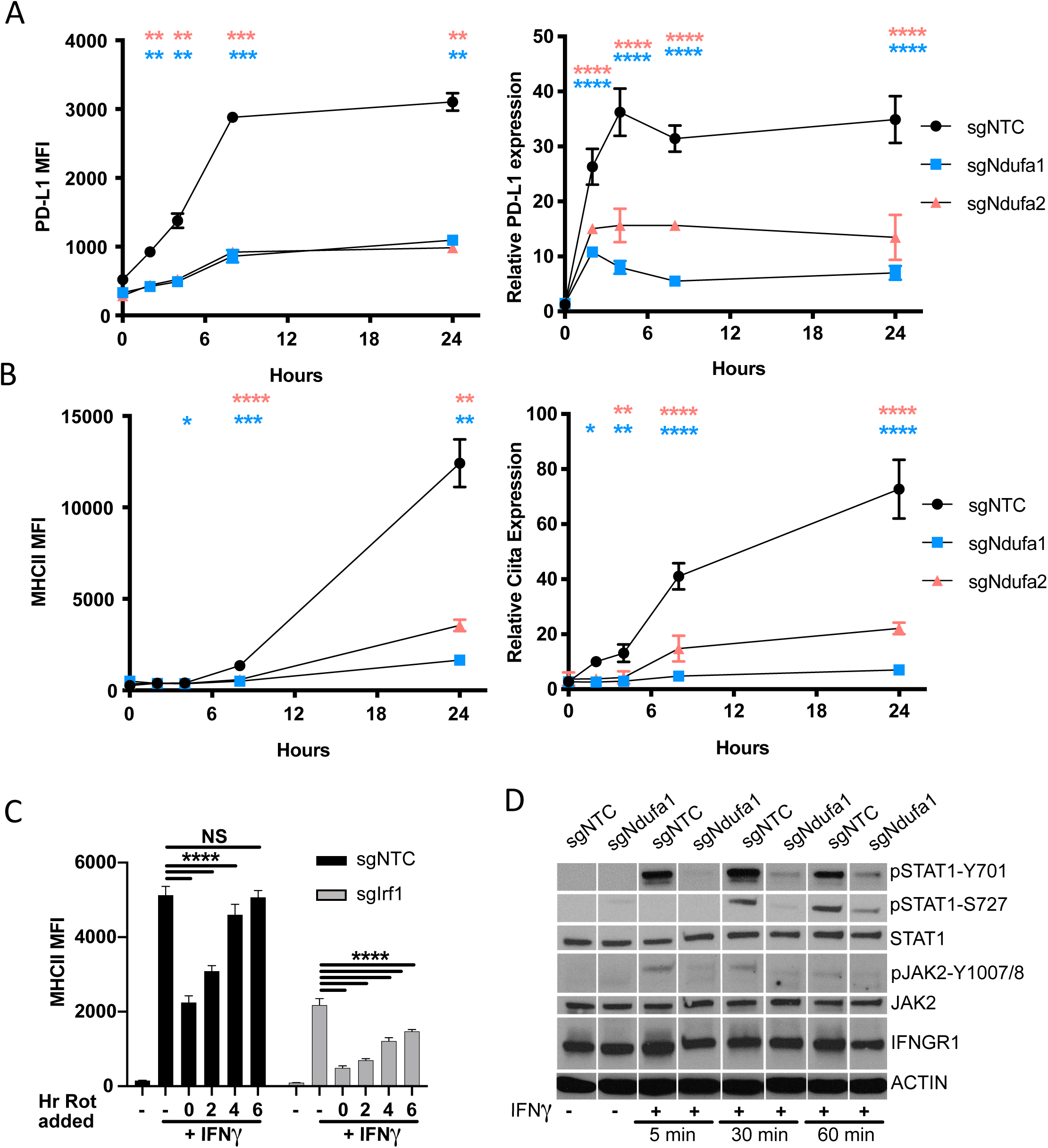
Complex I inhibition reduces IFNγ receptor activity. A) PD-L1 transcript was quantified by qRT-PCR using ΔΔCt relative to *β-Actin* in macrophages of the indicated genotype after stimulation with 10ng/mL IFNγ. PD-L1 MFI was determined at the same time points by flow cytometry. B) *Ciita* transcript was quantified by qRT-PCR using ΔΔCt relative to *β-Actin Gapdh* in macrophages of the indicated genotype after stimulation with 10ng/mL IFNγ. MHCII MFI was determined at the same time points by flow cytometry. Data shown are from biological triplicate samples with technical replicates for RT-PCR experiments and are representative of two independent experiments. C) sgNTC (left) or sg*Irf1* (right) macrophages were cultured for 24 hours with or without IFNγ stimulation. At 2 hour intervals post-IFNγ stimulation, rotenone was added. After 24 hours of stimulation, cells were harvested and surface expression of MHCII (MFI) was quantified by flow cytometry. Data are mean ± the standard deviation for 3 biological replicates and are representative of two independent experiments. Statistical testing was performed by one-way ANOVA with Tukey correction for multiple hypothesis testing. D) Control (NTC) or sg*Ndufa1* macrophages were stimulated with IFNγ for the indicated times, and cell lysates analyzed by immunoblot for STAT1 abundance and phosphorylation (Y701 and S727), JAK2 abundance and phosphorylation (Y1007/8), and IFNGR1. Beta-Actin was used as a loading control. Data are representative of three independent experiments. Results shown are from a single experiment analyzed on three parallel blots. p values of 0.05, 0.01, 0.001, and 0.001are indicated by *, **, ***, and ****.

IFNγ rapidly induces the transcription of a large number of STAT1 target genes, including IRF1, which amplifies the response. The relative impact of complex I inhibition on the immediate transcriptional response versus the subsequent IRF1-dependent amplification was initially assessed by altering the timing of complex I inhibition. As the addition of rotenone was delayed relative to IFNγ stimulation, the ultimate effect on MHCII expression was diminished (Figure 6C). If rotenone was added more than 4 hours after IFNγ, negligible inhibition was observed by 24 hours, indicating that early events were preferentially impacted by rotenone. To more formally test the role of IRF1, this study was performed in macrophages harboring a CRISPR-edited *Irf1* gene. While the level of MHCII induction was reduced in the absence of IRF1, the relative effect of rotenone addition over time was nearly identical in sg*Irf1* and sgNTC cells. Thus, mitochondrial function appeared to preferentially impact the initial transcriptional response to IFNγ upstream of IRF1.

Ligand induced assembly of the IFNGR1-IFNGR2 receptor complex results in the phosphorylation and transactivation of janus kinases 1 and 2 (JAK1, JAK2).

Autophosphorylation of JAK2 at tyrosine residues 1007/1008 positively regulates this cascade and serves as a marker of JAK2 activation. These activating events at the cytoplasmic domains of the IFNGR receptor complex facilitate STAT1 docking and phosphorylation at tyrsone-701 (Y701), a prerequisite for the IFNγ response. Additional STAT1 phosphorylation at serine-727 can amplify signaling. To determine if complex I is required for these early signal transduction events, we examined the activation kinetics by immunoblot (Figure 6D). The total abundances of IFNGR1, STAT1, and JAK2, were constant in sgNTC and sg*Ndufa1* cells in the presence and absence IFNγ-stimulation. While we detected robust phosphorylation of JAK2 Y1007/8, STAT1-Y701, and STAT1-S727 over time following IFNγ treatment in sgNTC cells, phosphorylation at all three sites was both delayed and reduced across the time-course in sg*Ndufa1* cells. We conclude that the loss of complex I function inhibits receptor proximal signal transduction events.

### Mitochondrial respiration in antigen presenting cells is required IFNγ-dependent T cell activation

As respiration affected both stimulatory and inhibitory antigen presenting cell (APC) functions, we sought to understand the ultimate effect of mitochondrial function on T cell activation. To this end, we generated myeloid progenitor cell lines from Cas9-expressing transgenic mice that can be used for genome-edited and differentiated into either macrophages or dendritic cells using M-CSF or FLT3L, respectively (110, 111). Macrophages differentiated from these myeloid progenitors demonstrated robust induction of all three markers that were the basis for the IFNγ stimulation screens (Supplementary Figure 2A-C). Further, both the IFNγ-mediated upregulation of these markers and the inhibitory effect of rotenone or oligomycin on their induction were indistinguishable from wild-type primary bone marrow-derived macrophages (Supplementary Figure 2D-F). In both macrophages and in dendritic cells (DCs), the induction of MHCII by IFNγ was inhibited by rotenone and oligomycin (Figure 7A). Unlike macrophages, murine DCs basally express MHCII and these inhibitors only repressed the further induction by IFNγ (Figure 7A,B).

**Figure 7.**
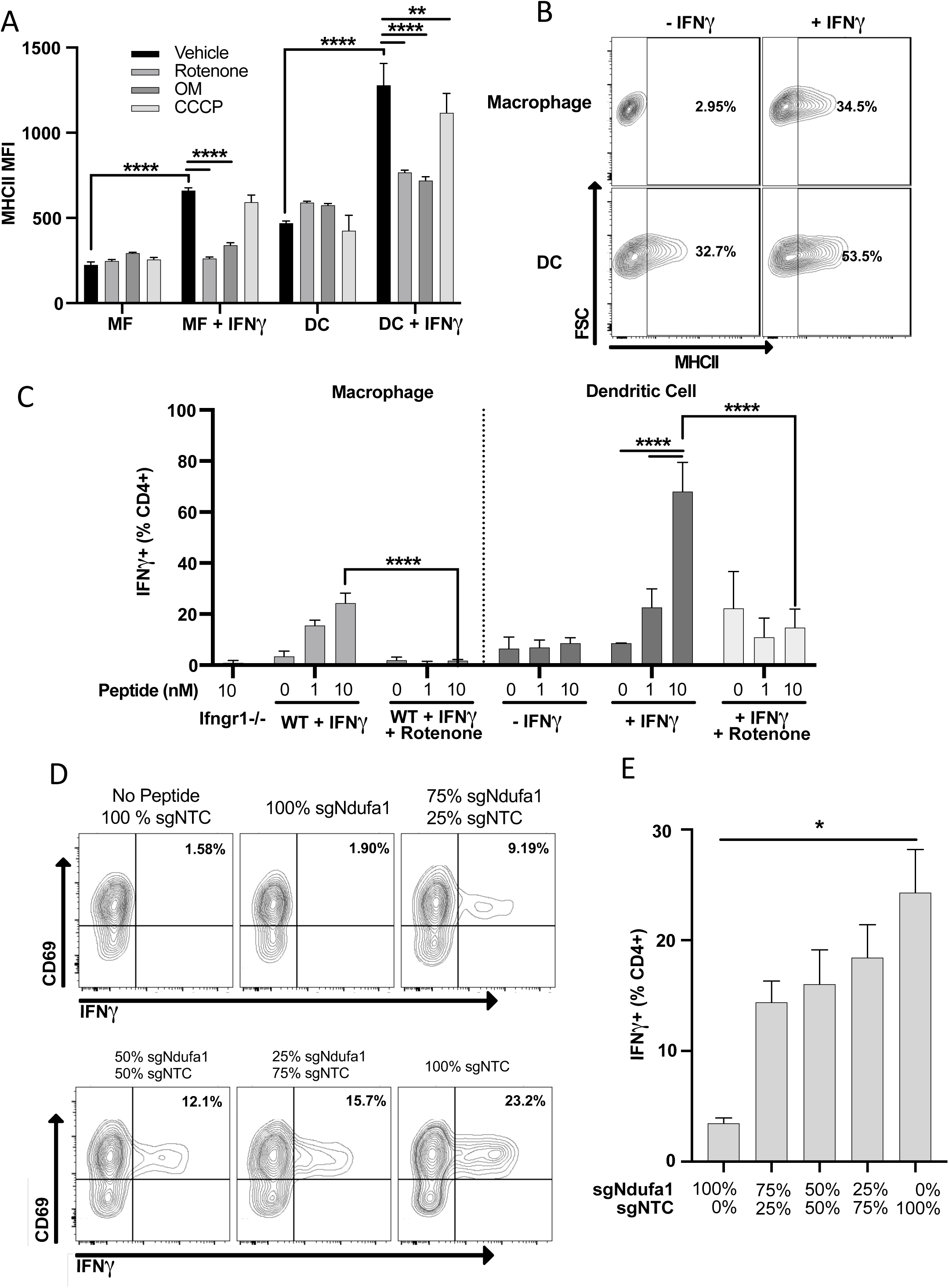
Mitochondrial respiration in antigen presenting cells is required IFNγ-dependent T cell activation. A) Cell surface expression of MHCII (MFI) in macrophages (MF) or dendritic cells (DC) derived from conditionally immortalized progenitor lines. IFNγ was added for 24 hours where indicated. Cells were treated with vehicle (DMSO), rotenone (10uM), oligomycin (OM, 2.5uM), or CCCP concurrent with IFNγ. Data are three biological replicates and are representative of at least two independent experiments. B) Contour plot of macrophage (top row) or dendritic cell (bottom row) MHCII expression in the absence of (left column) or following (right column) stimulation with IFNγ for 24 hours. Representative samples were selected from (A). The percent MHCII positive are indicated for each of the conditions. C) CD4+ T cell activation as measured by the percent of live cells positive for IFNγ by intracellular cytokine staining. Prior to co-culture with T cells, APCs were stimulated with the indicated combinations of IFNγ (10ng/mL), and/or rotenone (10uM) for 24 hours. After washing and pulsing with ESAT-61-15 at the indicated concentrations (nm.), T cells were added to APCs at an effector to target (E:T) ratio of 1:1, and co-cultured for a total of 5 hours. Data are representative of two independent experiments. Data are mean ± the standard deviation for 3 biological replicates. Statistical testing was performed by one-way ANOVA with Tukey correction for multiple hypothesis testing. D and E) sg*Ndufa1* or NTC macrophages were differentiated from immortalized progenitors, and mixed at the ratios indicated (labeled as percent of KO cells). Mixed cultures were stimulated with IFNγ for 24 hours, peptide loaded, and co-cultured with CD4+ T cells (E:T 1:1). Production of IFNγ was measured by ICS and quantified as the percent of cells positive for staining by flow cytometry. Representative contour plots (D) and quantification (E) of the experiment are shown. Data shown are for biological triplicate samples and are representative of two independent experiments. p values of 0.05, 0.01, 0.001, and 0.001are indicated by *, **, ***, and ****.

Both macrophages and DCs were used to determine if the inhibition of complex I in APCs reduces T cell activation. Both types of APCs were stimulated with IFNγ overnight with or without rotenone before washing cells to remove rotenone and ensure T cell metabolism was unperturbed. APCs were then pulsed with a peptide derived from the *Mycobacterium tuberculosis* protein ESAT-6, and co-cultured with ESAT-6-specific CD4+ T cells from a TCR transgenic mouse (112). T cell activation was assayed by intracellular cytokine staining for IFNγ. In macrophages, T cell stimulation relied on pretreatment of the APC with IFNγ, as a macrophage line lacking the *Ifngr1* gene was unable to support T cell activation. Similarly, inhibition of complex I in macrophages completely abolished antigen-specific T cell stimulation (Figure 7C). DCs did not absolutely require IFNγ pretreatment to stimulate T cells, likely due to the basal expression of MHCII by these cells. Regardless, rotenone treatment of DC abrogated the IFNγ-dependent increase in T cell stimulation (Figure 7C).

To confirm the effects of complex I inhibition on T cell activation using a genetic approach and confirm that complex I inhibition acted in a cell-autonomous mechanism, we generated *Ndufa1* knockout myeloid progenitors (Hox-sg*Ndufa1*). Following differentiation into macrophages, Hox-sg*Ndufa1* demonstrated glycolytic dependence and the inability to generate ATP by OXPHOS compared to control Hox-sgNTC macrophages (Supplementary Figure 2G). Having confirmed the expected metabolic effects of Ndufa1 loss, Hox-sg*Ndufa1* and Hox-sgNTC macrophages were mixed at various ratios. Mixed cultures were then stimulated with IFNγ, peptide pulsed, and co-cultured with antigen-specific CD4+ T cells. In agreement with our chemical inhibition studies, we found strong correlation between complex I activity in the APC population and T cell stimulatory activity (Figure 7D-E). Together, these data confirm that the IFNγ-dependent augmentation of T cell stimulatory activity depends on complex I function in both macrophages and DCs.

## Discussion

IFNγ-mediated control of APC function is central to shaping a protective immune response, and the canonical IFNγ signal transduction pathway has been elucidated in exquisite detail (113). Our study demonstrates that unbiased genetic analyses can reveal a multitude of unexpected cellular regulators, even for a well-characterized process such as IFNγ signaling. By independently assessing genetic determinants of stimulatory and inhibitory molecule expression, we discovered mechanisms of regulation that preferentially affect the induction of different cell surface proteins. These results begin to explain how a single cytokine can induce functionally distinct downstream responses in different contexts. These data also suggest new strategies to modulate individual co-receptors to either stimulate or inhibit T cell activation. Another strength of our parallel screen approach was the increased power to identify shared mechanisms that control IFNγ-mediated regulation across all screens. Our pooled analysis identified mitochondrial respiration, and in particular complex I, as essential for IFNγ-responses in APCs. We determined that complex I is required for the IFNγ-mediated induction of key immune molecules and is necessary for antigen presentation and T cell activation. These findings uncover a new dependency between cellular metabolism and the immune response.

Our genetic and chemical inhibition data demonstrated that mitochondrial respiration is necessary for early events in signal transduction from the IFNγ receptor complex, and complex I of the respiratory chain is specifically required. While IFNγ stimulation has been reported to mediate a reduction in oxygen consumption and a shift to aerobic glycolysis over time (66), the requirement of mitochondrial respiration in IFNγ responses has not been assessed previously. Our results indicate that complex I is required for IFNγ signaling regardless of these metabolic shifts. Complex I is a metabolic hub with several core functions that cumulatively recycle nicotinamide adenine dinucleotide (NAD+), reduce ubiquinol, and initiate the PMF for ATP generation. While any or all of these physiologic processes could contribute to IFNγ signaling, the differential effects of chemical inhibitors narrow the possibilities. Both rotenone and oligomycin inhibit the IFNγ response, and block electron flux through complex I either directly or indirectly. In contrast, the ionophore CCCP disrupts the PMF and ATP generation without inhibiting electron transfer, and does not affect IFNγ signaling. These data indicate that the reduction state of the quinone pool and ATP generation do not regulate IFNγ responses in our system. Instead, complex I-dependent regeneration of NAD+ is the most likely regulator of IFNγ signaling. Indeed, NAD+ synthesis via either the *de novo* or salvage pathway is necessary for a variety of macrophage functions (114–116). Very recent work demonstrates an important role for NAD+ in STAT1 activation and PD-L1 induction by IFNγ in hepatocellular carcinoma cells (117). In this setting, inhibition of NAD+ synthesis reduces the abundance of phospho-STAT1 by disrupting a direct interaction with the Ten-eleven translocation methylcytosine dioxygenase 1 (TET1). It remains unclear if a similar interaction occurs in the myeloid cells that are the focus of our work, as TET1 is expressed at very low levels in macrophages and splenic DC (118). Regardless, these observations indicate that both NAD+ synthesis and its regeneration via mitochondrial respiration contribute to the IFNγ response in diverse cell types. This recently revealed interaction between metabolism and immunity could contribute to the observed association between NAD+ homeostasis and inflammatory diseases (116), as well as the efficacy of checkpoint inhibitor therapy for cancer (117).

In the APC setting, we found that T cell activation required mitochondrial respiration. While complex I function, MHCII and CD40 expression all largely correlate with T cell stimulation, our data indicate that additional IFNγ-inducible pathways also contribute to this activity. For example, unstimulated DCs basally express similar levels of MHCII as IFNγ- stimulated macrophages but are unable to productively present antigen to T cells. This observation suggests that additional aspects of antigen processing, presentation, or co-stimulation are IFNγ- and complex I-dependent. Similarly, MHCI presentation machinery is transcriptionally induced upon IFNγ stimulation (7, 119) and the induction of molecules recognized by donor unrestricted T cells, such as MR1 and CD1, might also require additional signals to function. The specific effects of mitochondrial respiration on the type and quality of the T cell response will depend on how these diverse antigen-presenting and co-signaling molecules are influenced by cellular metabolic state.

The observation that IFNγ signaling depends on mitochondrial respiration provides a stark contrast to the well-established glycolytic dependency of many phagocyte functions, such as TLR signaling. This metabolic dichotomy between proinflammatory TLR signals and the IFNγ response mirrors known regulatory interactions between these pathways. For example, TLR stimulation has been shown to inhibit subsequent IFNγ responses, via a number of target gene-specific mechanisms (120–124). However, TLR stimulation also results in the disassembly of the ETC (123, 124), which our observations predict to inhibit STAT1 phosphorylation and IFNγ signaling at the level of the receptor complex. More generally, our work suggests fundamental metabolic programs contribute to the integration of activation signals by APC and influence the ultimate priming of an immune response.

## Materials and Methods

### Cell culture

Cells were cultured in Dulbecco’s Modified Eagle Medium (Gibco 11965118) supplemented with 10% fetal bovine serum (Sigma F4135), sodium pyruvate (Gibco 11360119), and HEPES (15630080). Primary bone marrow-derived macrophages (BMDMs) were generated by culturing bone marrow in the presence of media supplemented with 20% L929 supernatant for 7 days.

Immortalized macrophage cell lines in C57Bl6/J and Cas9-EGFP were established in using J2 retrovirus from supernatant of CREJ2 cells as previously described(125). Briefly, isolated bone marrow was cultured in the presence of media enriched with 20% L929 supernatant. On day 3, Cells were transduced with virus and cultured with virus for 2 days. Over the next 8 weeks, L929 media was gradually reduced to establish growth factor independence.

Conditionally immortalized myeloid progenitor cell lines were generated by retroviral transduction using an estrogen-dependent Hoxb8 transgene as previously described(110). Briefly, mononuclear cells were purified from murine bone marrow using Ficoll-Paque Plus (GE Healthcare 17144002) and cultured in RPMI (Gibco 11875119) containing 10% fetal bovine serum (Sigma F4135), sodium pyruvate (Gibco 11360119), and HEPES (15630080), IL-6 (10ng/mL; Peprotech #216-16), IL-3 (10ng/mL; Peprotech #213-13), and SCF (10ng/mL; Peprotech #250-03) for 48 hours. Non-adherent bone marrow cells from C57Bl6/J (Jax 000664), Cas9-EGFP knockin (Jax 026179), or Ifngr1 knockout (Jax 003288) mice were transduced with ER-Hoxb8 retrovirus. After transduction cells were cultured in with media supplemented with supernatant from B16 cells expressing GM-CSF and 10uM estradiol (Sigma E8875) to generate macrophage progenitor cell lines or in media supplemented with supernatant from B16 cells expressing FLT3L and 10uM estradiol (Sigma E8875) to generate dendritic cell progenitor lines. To differentiate macrophages, progenitors were harvested and washed twice with PBS to remove residual estradiol and cultured in L929 supplemented media as above. To differentiate dendritic cells(111), progenitors were harvested, washed 2x with PBS, and cultured in FLT3-enriched complete RPMI for 8-10 days.

Human monocyte-derived macrophages (MDM) were differentiated from mononuclear cells of healthy donors. Briefly, peripheral blood mononuclear cells (PBMCs) were isolated from whole blood using Ficoll-Paque-PLUS (GE Healthcare 17144002). CD14+ monocytes were purified using MojoSort™ Human CD14 Nanobeads (Biolegend 480093) according to the manufacturer’s protocol. Cells were cultured in RPMI with 10% FBS, sodium pyruvate, and HEPES and supplemented with recombinant GM-CSF (50ng/mL, Peprotech 300-03) for 6 days. Thaws were harvested using Accutase (Gibco A1110501).

### Cell stimulations

Murine IFNγ (Peprotech 315-05) and human IFNγ (Peprotech 300-02) were used at 10ng/mL unless otherwise indicated in the figure legends. Murine TNF (315-01A) was used at 25ng/mL. Pam3CSK4 (Invivogen tlrl-pms) was used at 200ng/mL.

### CRISPR screens

A clonal macrophage cell line stably expressing Cas9 (L3) was established as described elsewhere(79). A plasmid library of sgRNAs targeting all protein coding genes in the mouse genome (Brie Knockout library, Addgene 73633) was packaged into lentivirus using HEK293T cells. HEK293T supernatants were collected and clarified, and virus was titered by quantitative real-time PCR and by colony counting after transduction of NIH3T3. L3 cells were transduced at a multiplicity of infection (MOI) of ∼0.2 and selected with puromycin 48 hours after transduction (2.5ug/mL). The library was minimally expanded to avoid skewing mutant representation and then frozen in aliquots in freezing media (90% FBS 10% DMSO).

Two replicate screens for MHCII, CD40, and PD-L1 were performed as follows: 2e8 cells of the knockout (KO) library was stimulated with IFNγ (10ng/mL; Peprotech 315-05) for 24 hours after which cells were harvested by scraping to ensure integrity of cell surface proteins. Cell were stained with TruStain FcX anti-mouse CD16/32 (Biolegend 101319) and LIVE/DEAD Fixable Aqua (Invitrogen L34957) per the manufacturer’s instructions. For each of the respective screens, stimulated library was stained for its respective marker with the following antibody: MHCII (APC anti-mouse I-A/I-E Antibody, Clone M5/114.15.2 Biolegend 107613), CD40 (APC anti-mouse CD40 Antibody, Clone 3/23 Biolegend 124611), or PD-L1 (APC anti-mouse CD274 (B7-H1, PD-L1) Antibody, Clone 10F.9G2 Biolegend 124311). Each antibody was titrated for optimal staining using the isogenic L3 macrophage cell line. Following staining, cells were fixed in 4% paraformaldehyde. High and low expressing populations were isolated by fluorescence activated cell sorting (FACS) using a BD FACS Aria II Cell Sorter. Bin size was guided by control cells which were unstimulated and to ensure sufficient library coverage (>25x unselected library, or >2e6 cells per bin). Following isolation of sorted populations, paraformaldehyde crosslinks were reversed by incubation in proteinase K (Qiagen) at 55 degrees for 6-8 hours. Subsequently, genomic DNA was isolated using DNeasy Blood and Tissue Kit (Qiagen 69504) according to the manufacturer’s instructions. Amplification of sgRNAs by PCR was performed as previously described(74, 126) using Illumina compatible primers from IDT, and amplicons were sequenced on an Illumina NextSeq500. Sequence reads were trimmed to remove adapter sequence and to adjust for staggered forward (p5) primer using Cutadapt v2.9. Raw sgRNA counts for each sorted and unsorted (input library) population was quantified using bowtie2 via MAGeCK to map reads to the sgRNA library index (no mismatch allowed); a sgRNAindex was modified to reflect genes transcribed by our macrophage cell line either basally or upon stimulation with IFNγ as previously published(79). Counts for sgRNAs were median normalized to account for variable sequencing depth.

### MAGeCK-MLE

We used MAGeCK-MLE to test for gene enrichment. Two separate analyses were performed in order to: (1) identify regulators of the IFNγ response, and (2) identify specific regulators of each of the screen targets. For both analyses, the baseline samples were the input libraries from each of the replicate screens in order to account for slight variabilities in library distribution for each screen. For (1), the generalized linear model was based on a design matrix that was “marker-blind” and only considered the bin of origin (i.e. MHCII-low, CD40-low, PD-L1-low v. MHCII-high, CD40-high, PD-L1-high). For (2), the design matrix was “marker-aware and bin-specific” to test for marker-specific differences (i.e. MHCII-low v. CD40-low v. PD-L1-low); the analysis was performed separately for each bin, low or high expressing mutants, to identify marker-specific positive and negative regulators, respectively. For each analysis, β scores (selection coefficient) for each gene were summed across conditions to allow for simultaneous assessment of positive and negative regulators across conditions. Data are provided in Supplementary Tables.

Gene-set enrichment analysis (GSEA) was performed using a ranked gene list as calculated from MAGeCK-MLE beta scores and false discovery rate (FDR). To facilitate the identification of positively and negatively enriched gene sets from the high and low expressing populations, the positive (“pos | beta”) and negative (“neg | beta”) beta scores for each gene were summed as described above (“beta_sum”). To generate a ranked gene list for GSEA, we employed Stouffer’s method to sum positive (“pos | z”) and negative (“neg | z”) selection z-scores, which were used to re-calculate p-values (“p_sum”) as has been previously described (127–129). Using these summative metrics, we calculated a gene score as: log10(p_sum) * (beta_sum). Genes were ranked in descending order and GSEA was performed with standard settings including “weighted” enrichment statistic and “meandiv” normalization mode. Analysis was inclusive of gene sets comprising of 10-500 genes that were compiled and made available online by the Bader lab (130, 131).

### Plasmids and sgRNA cloning

Lentivirus was generated using HEK293T cells using packaging vector psPAX2 (Addgene#12260) and envelope plasmid encoding VSV-G. Transfections used TransIT-293 (MirusBio MIR 2704) and plasmid ratios according to the manufacturer’s instructions. For the generation of retrovirus, pCL-Eco in place of separate packaging and envelope plasmid. Retrovirus encoding the ER-Hoxb8 transgene was kindly provided by David Sykes.

sgOpti was a gift from Eric Lander & David Sabatini (Addgene plasmid #85681)(132). Individual sgRNAs were cloned as previously described. Briefly, annealed oligos containing the sgRNA targeting sequence were phosphorylated and cloned into a dephosphorylated and BsmBI (New England Biolabs) digested SgOpti (Addgene#85681) which contains a modified sgRNA scaffold for improved sgRNA-Cas9 complexing. Use of sgOpti derivatives for delivery of multiple sgRNAs was performed as detailed elsewhere(79). The sgRNA targeting sequences used for cloning were as follows:

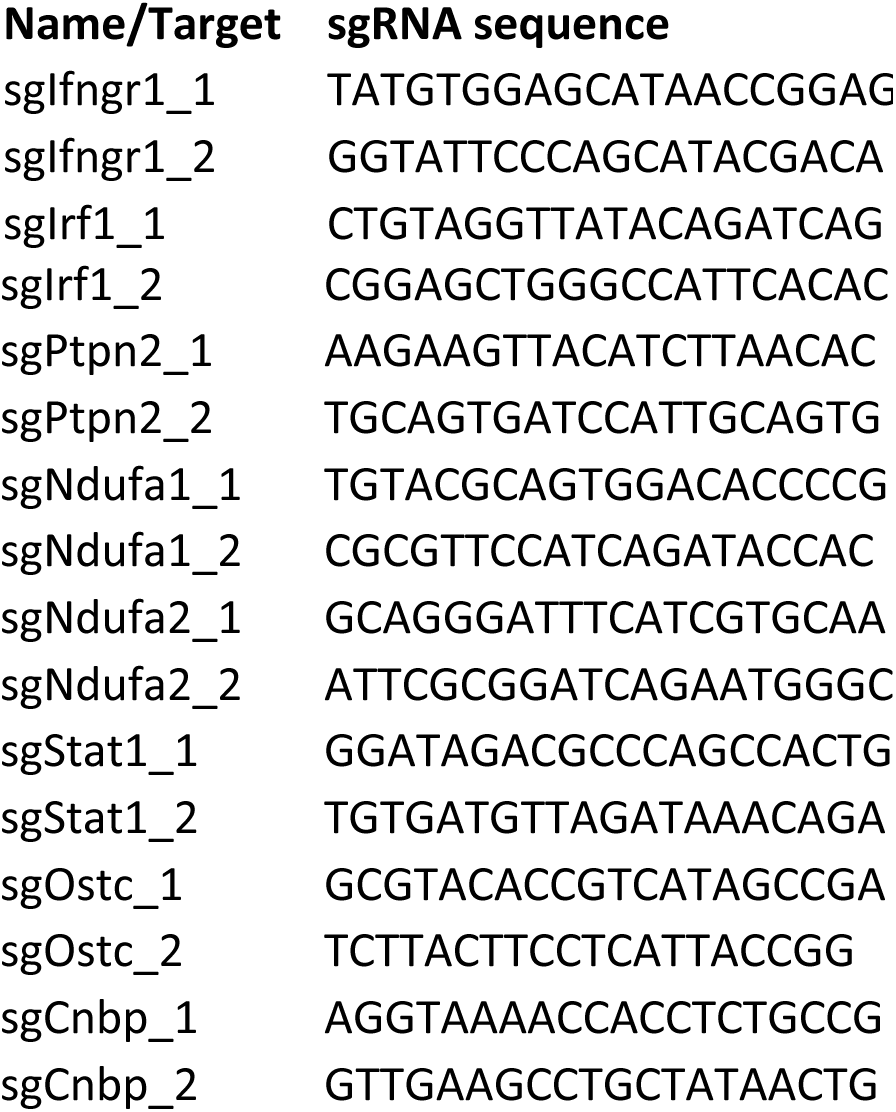

### Flow cytometry

Cells were harvested at the indicated times post-IFNγ stimulation by scrapping to ensure intact surface proteins. Cells were pelleted and washed with PBS before staining with TruStain FcX anti-mouse CD16/32 (Biolegend 101319) or TruStain FcX anti-human (Biolegend 422301) and LIVE/DEAD Fixable Aqua (Invitrogen L34957) per the manufacturer’s instructions. The following antibodies were used as indicated in the figure legends:

APC-Fire750 anti-mouse I-A/I-E Antibody, Clone M5/114.15.2 Biolegend 107651

PE anti-mouse CD40 Antibody, Clone 3/23 Biolegend 124609

Brilliant Violet 421™ anti-mouse CD274 (B7-H1, PD-L1) Antibody, Clone 10F.9G2 Biolegend 124315

Alexa Fluor® 647 anti-human CD54 Antibody, Clone HCD54, Biolegned 322718

PE anti-human CD40 Antibody, Clone 5C3, Biolegned 334307

Brilliant Violet 421™ anti-human CD274 (B7-H1, PD-L1) Antibody, Clone 29E.2A3, Biolegend 329713

APC/Fire™ 750 anti-human HLA-DR Antibody, Clone L243, Biolegend 307657

For intracellular cytokine staining, cells were treated with brefeldin A (Biolegend 420601) for 5 hours before harvesting. Following staining and fixation, cells were permeabilized (Biolegend 421002) and stained according to the manufacturer’s protocol using the following antibodies: PE anti-mouse IFN-γ Antibody, Biolegend 505807

Surface protein expression was analyzed on either a MacsQuant Analyzer or Cytek Aurora. All flow cytometry analysis was done in FlowJo V10 (TreeStar).

### Chemical inhibitors

All chemical inhibitors were used for the duration of cell stimulation unless otherwise stated. Rotenone (Sigma R8875) was resuspended in DMSO and used at 10uM unless indicated otherwise in the figure legend. Oligomycin (Cayman 11342) was resuspended in DMSO and used at 2.5uM unless otherwise indicated. CCCP (Cayman 25458) was resuspended in DMSO and used at 1.5uM unless indicated otherwise. 1400W hydrochloride (Cayman 81520) was resuspended in culture media, filter sterilized and used immediately at 25uM unless otherwise indicated. N-acetyl-L-Cysteine (NAC, Cayman 20261) was resuspended in culture media, filter sterilized and used immediately at 10mM. DMOG (Cayman 71210) was resuspended in DMSO and used at 200uM. UK5099 (Cayman 16980) was resuspended in DMSO and used at 20uM. 2-deoxy-D-Glucose (2DG, Cayman 14325) was resuspended in culture media, filter sterilized and used at 1mM or at the indicated concentrations immediately. MitoTEMPO hydrate (Cayman 16621) was resuspended in DMSO and used at the indicated concentrations.

For experiments that used defined minimal media with carbon supplementation, D-galactose, sodium pyruvate, and D-glucose were used at 10mM in DMEM without any carbon (Gibco A1443001). For establishment of macrophage cell line with diminished mitochondrial mass, cells were continuously cultured in linezolid (LZD) (Kind gift from Clifton Barry) for 4 weeks at 50 μg/mL or DMSO control. Both LZD-conditioned and DMSO control lines were supplemented with uridine at 50 μg/mL. Prior to experimentation, cells were washed with PBS and cultured without linezolid for at least 12 hours.

### ELISA and nitric oxide quantification

The following kits were purchased from R and D Systems or Biolegend for quantifying protein for cell supernatants:

Mouse IL-6 DuoSet ELISA (DY406) or Biolegend ELISAmax (431301)

Mouse TNF-alpha DuoSet ELISA (DY410) or Biolegend ELISAmax (430901)

Mouse IFN-gamma DuoSet ELISA (DY485)

Human IL-1 beta/IL-1F2 DuoSet ELISA (DY201)

Human TNF-alpha DuoSet ELISA (DY210)

Nitric oxide was quantified from cell supernatants using the Griess Reagent System according to the manufacturer’s instructions (Promega G2930). For these experiments, cell culture media without phenol red (Gibco A1443001 or Gibco 31053028).

### RNA isolation and quantitative real-time PCR

To isolate RNA, cells were lysed in TRIzol (15596026) according to manufacturer’s instructions. Chloroform was added to lysis at ratio of 200uL chloroform per 1mL TRIzol and centrifuged at 12,000 x g for 20 minutes at 4C. The aqueous layer was separated and added to equal volume of 100% ethanol. RNA was isolated using the Zymo Research Direct-zol RNA extraction kit. Quantity and purity of the RNA was checked using a NanoDrop and diluted to 5ng/uL in nuclease-free water before use. Quantitative real-time PCR was performed using NEB Luna® Universal One-Step RT-qPCR Kit (E3005) or the Quantitect SYBR green RT-PCR kit (204243) according to the manufacturer’s protocol and run on a Viia7 thermocycler or StepOne Plus Theromocycler. Relative gene expression was determined with ddCT method with beta-Actin transcript as the reference.

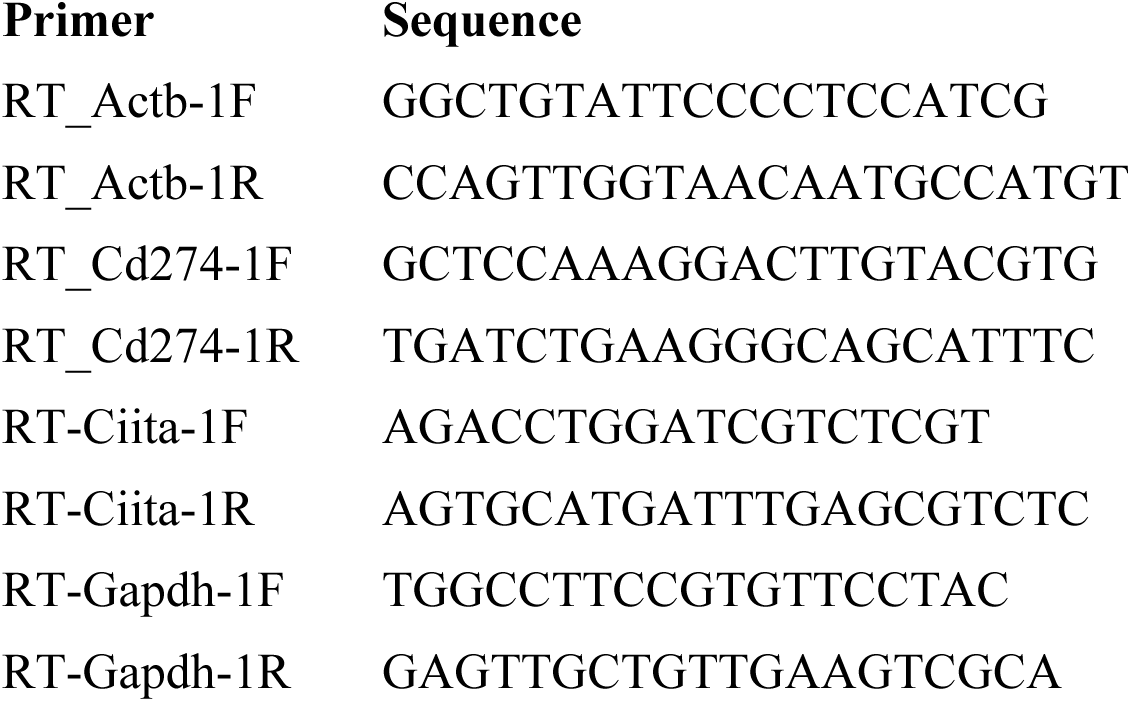

### Quantification of mitochondrial genomes

Genomic DNA was isolated from cell pellets using the DNeasy Blood and Tissue Kit (Qiagen 69504). Quantitative PCR was run using NEB Luna® Universal One-Step RT-qPCR without the RT enzyme mix and run on a Viia7 thermocycler. Relative quantification of mitochondrial genomes was determined by measuring the relative abundance of mitochondrially encoded gene Nd1 to the abundance of nuclear encoded Hk2 as has been described elsewhere(133). All primers are detailed in attached table.

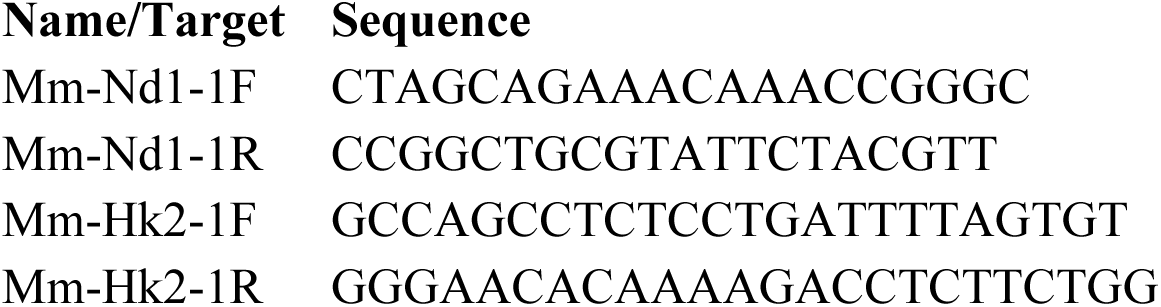

### Immunoblot

At the indicated times following stimulation, cells were washed with PBS once and lysed in on ice using the following buffer: 1% Triton X-100, 150mM NaCl, 5mM KCl, 2mM MgCl2, 1mM EDTA, 0.1% SDS, 0.5% DOC, 25mM Tris-HCl, pH 7.4, with protease and phosphatase inhibitor (Sigma #11873580001 and Sigma P5726). Lysates were further homogenized using a 25g needle and cleared by centrifugation before quantification (Pierce™ BCA Protein Assay Kit, 23225).

Parallel blots were run with the same samples, 15ug per well. The following antibodies were used according to the manufacturer’s instructions:

Purified anti-STAT1 Antibody Biolegend Clone A15158C

Purified anti-STAT1 Phospho (Ser727) Antibody, Biolegend Clone A15158B

Phospho-Stat1 (Tyr701) Rabbit mAb, Cell Signaling Technology Clone 58D6

Jak2 XP® Rabbit mAb, Cell Signaling Technology Clone D2E12

Phospho-Jak2 (Tyr1007/1008) Antibody, Cell Signaling Technology #3771S

Anti-mouse β-Actin Antibody, Santa Cruz Biotechnology Clone C4

Biotin anti-mouse CD119 (IFN-γ R α chain) Antibody, Biolegend Clone 2E2

Goat anti-Rabbit IgG (H+L) Secondary Antibody, HRP, Invitrogen 31460

Goat anti-Mouse IgG (H+L) Secondary Antibody, HRP, Invitrogen 31430

HRP-Conjugated Streptavidin, Thermo Scientific N100

### Bioenergetics Assay

Relative glycolytic and respiratory capacity were determined as has previously been demonstrated(134). Briefly, cellular ATP levels were determined using CellTiter-Glo® 2.0 Cell Viability Assay (Promega G9241) according to the manufacturer’s protocol. Cells were grown in the conditions indicated in the figure legends for 4 hours unless stated otherwise. ATP levels were normalized according to the figure legend.

### T cell activation assay

We used a previously established co-culture system to assess antigen presentation to Ag-specific T cells. Briefly, C7 CD4+ T cells were isolated from transgenic C7 mice, respectively and stimulated in vitro with irradiated splenocytes pulsed with the ESAT-61-15 peptide, in complete media (RPMI with 10% FBS) containing IL-2 and IL-7. After the initial stimulation, the T cells were split every two days for 3-4 divisions and rested for two to three weeks. After the initial stimulation, the cells were cultured in complete media containing IL-2 and IL-7. The following synthetic peptide epitopes were used as antigens from New England Peptide (Gardener, MA): ESAT-61-15 (MTEQQWNFAGIEAAA).

For use in co-culture assay, T cells were added to peptide-pulsed macrophages as described in figure legends at an effector to target ratio of 1:1. Following 1 hours of co-culture, brefeldin A was added for 5 hours before assessing intracellular cytokine production by ICS.

### Quantification of subunit effects on N-module

We used publicly available proteomics data in which the protein abundance of all complex I subunit was measured when each subunit was genetically deleted(93). As determined empirically by the authors, the N-module components included: NDUFA1, NDUFA2, NDUFS1, NDUFV2, NDUFA6, NDUFS6, NDUFA7, NDUFS4, and NDUFV3. The relative effect of each subunit (using a knockout of that subunit) on N-module protein stability was calculated as the sum of the median log2 ratio of each of the above mentioned subunits, minus the median log2 ratio of itself (since it is knocked out).

### Statistical Analysis and Figures

Statistical analysis was done using Prism Version 8 (GraphPad) as indicated in the figure legends. Data are presented, unless otherwise indicated, as the mean +/- the standard deviation. Figures were created in Prism V8 or R (Version 3.6.2). MAGeCK-MLE was used as part of MAGeCK-FLUTE package v1.8.0.

## Acknowledgements

We thank all the members of the Sassetti, Behar, and Olive labs for critical feedback and input throughout the project. A special thank you to Megan K. Proulx, Mario Meza Segura and the donors for their assistance and expertise to the human macrophage derivation We thank the flow cytometry core at UMMS for their help in all experiments requiring flow cytometry. This work was supported by startup funding to AJO provided by Michigan State University, support from the Arnold O. Beckman Postdoctoral fellowship to AJO and grants from the NIH (AI146504, AI132130), DOD (W81XWH2010147), and USDA (NIFA HATCH 1019371). All data is being deposited into the appropriate databases and is available upon request.

## Supplementary Figure Legends

**Figure S1. Related to Figure 3.**
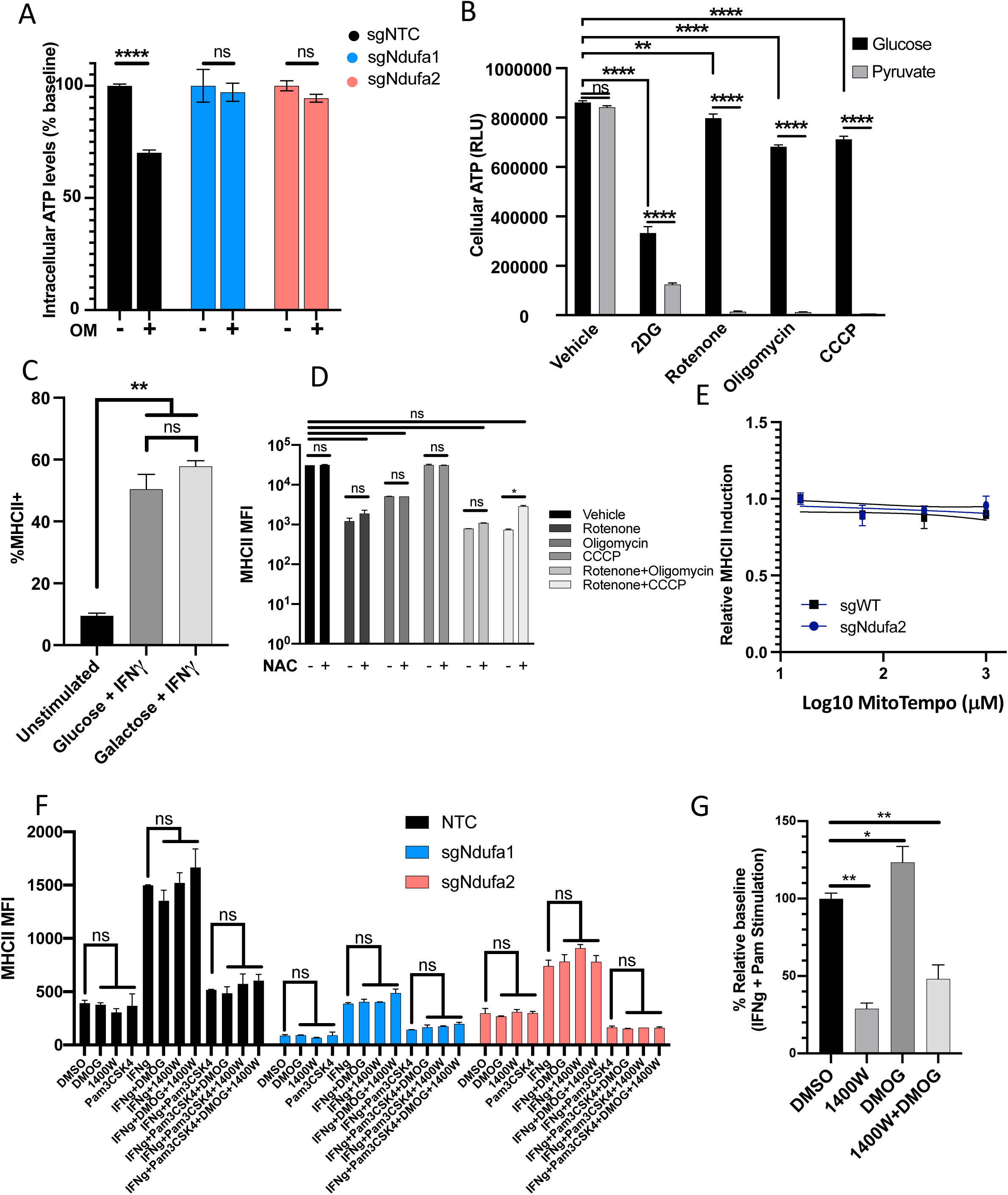
**A)** sgNTC, sg*Ndufa1*, sg*Ndufa2* cells cultured in complete media and treated with or without oligomycin (2.5μM) for 4 hours. Relative ATP levels were determined as in Figure 2A **B)** Intracellular ATP levels quantified as relative light units (RLU) using CellTiterGlo2.0 (Promega) for macrophages in specified growth conditions for 4 hours. Concentrations of carbon source and inhibitors are indicated in Materials and Methods. **C)** Macrophages were cultured in either glucose or galactose and stimulated with IFNγ for 24 hours. Following stimulation, the proportion of cells with MHCII expression was determined by flow cytometry. **D)** Macrophages were cultured in conditions as described in Figure 4H. For each condition, cells were stimulated with IFNγ or IFNγ and N-acetylcysteine (NAC) for 24 hours after which cell surface levels of MHCII were quantified. **E)** Control or complex I mutant (sg*Ndufa2*) macrophages were stimulated with IFNγ for 24 hours with increasing doses of mitochondrial reactive oxygen species scavenger MitoTempo. For each concentration, values are plotted as a fold change relative to no scavenger; Mean ± the standard deviation for 2 biological replicates of each condition. **F)** Control or complex I deficient macrophages were stimulated with IFNγ for 24 hours with or without the addition of DMOG or 1400W. Following stimulation, the proportion of cells with MHCII expression was determined by flow cytometry. G) Nitric oxide was measured using Griess Reagent System (Promega) from cell supernatants following stimulation with IFNγ and Pam3CSK4 for 24 hours with or without the addition of DMOG or 1400W. Relative nitric oxide levels were calculated as a percent relative to control (IFNγ and Pam3CSK4 with DMSO). All data are representative of at least two independent experiments. Statistical testing was performed using one-way ANOVA with Holm-Sidak multiple comparison correction. p values of 0.05, 0.01, 0.001, and 0.001are indicated by *, **, ***, and ****.

**Figure S2. Related to Figure 7.**
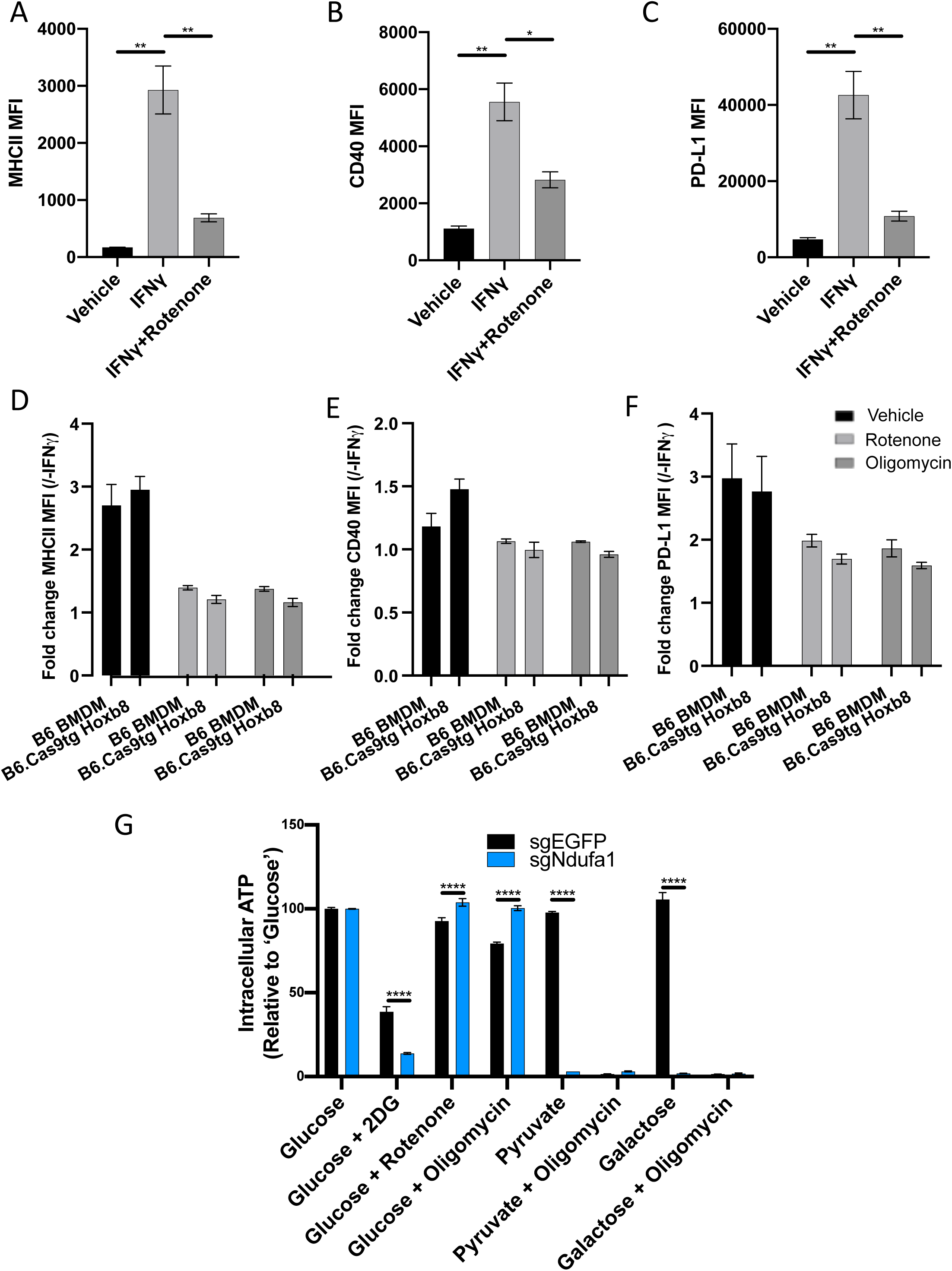
A-C) Myeloid progenitors cells were conditionally immortalized by transducing murine bone marrow with an estrogen-dependent Hoxb8 transgene which maintains stem-like properties. Following differentiation of progenitors into macrophages using M-CSF enriched conditioned media, macrophages were stimulated with IFNγ with or without rotenone. 24 hours after stimulation, cell surface levels of (A) MHCII, (B) CD40, (C) and PD-L1 were quantified by flow cytometry. Data are representative of 3 independent experiments and are the mean ± the standard deviation for 2 biological replicates. Statistical testing was performed by one-way ANOVA with Tukey correction for multiple hypothesis testing. D-F) As in panels A-C, macrophages from either immortalized macrophage progenitors or primary bone marrow were stimulated with IFNγ with or without rotenone or oligomycin. 24 hours after stimulation, cell surface levels of (D) MHCII, (E) CD40, (F) and PD-L1 were quantified by flow cytometry. G). Wild-type or ΔNdufa1 macrophages derived from Hoxb8-immortalized bone marrow progenitors were cultured in the specified media and inhibitor condition before total intracellular ATP was quantified by CellTiterGlo2.0. For each genotype, values are relative to “glucose” control. Mean ± the standard deviation for 2 biological replicates of each condition. p values of 0.05, 0.01, 0.001, and 0.001are indicated by *, **, ***, and ****.

